## Supplemental File 1 for "A SMARTTR workflow for multi-ensemble atlas mapping and brain-wide network analysis"

### SUPPLEMENTAL FIGURES AND FIGURE LEGENDS

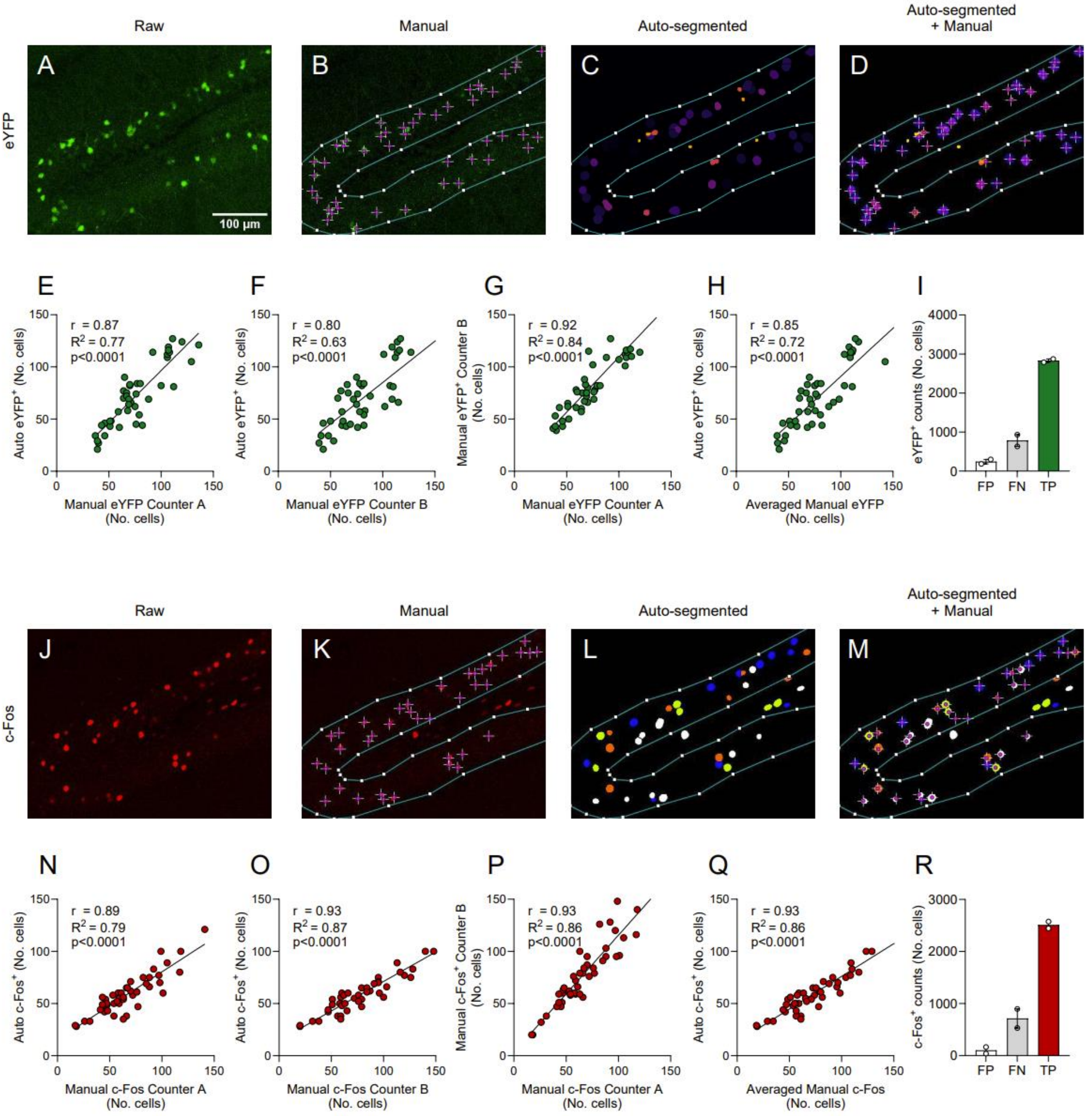


**Figure S1. Automated segmentation yields comparable results to manual cell counting. (A)** A representative image of the raw eYFP immunolabelled signal (channel 1) in the dentate gyrus (DG). **(B)** An ROI was drawn around the granule cell layer of the dentate gyrus and $eYFP^+$ cells within the boundaries of the ROI were manually counted. **(C)** Detected 3D objects representing $eYFP^+$ cells that were automatically segmented from the raw representative image. **(D)** Qualitative overlay of the manual $eYFP^+$ cell counts and the auto-segmented objects. **(E)** A high correlation (R = 0.87) was found between manual $eYFP^+$ cell counts by

Counter A and the number of auto-segmented objects. Each dot represents one ROI counted. (**F**) A high correlation (R = 0.80) was found between manual $eYFP^+$ cell counts by Counter B and the number of auto-segmented objects. **(G)** There was high inter-rater correlation (R = 0.92) between manual $eYFP^+$ cell counts. **(H)** Averaged manual $eYFP^+$ cell counts showed overall high correlation (R = 0.85) with auto-segmented counts. **(I)** Counts of all false positive (FP), false negative (FN), and true positive (TP) cell counts when comparing auto-segmented $eYFP^+$ counts and manual counts. **(J)** Representative image of the raw c-Fos immunolabelling signal (channel 2) in the DG. **(K)** Manual c-$Fos^+$ cell counts within the drawn ROI. **(L)** Automatically segmented 3D objects representing c-$Fos^+$ cells. **(M)** Overlay of manual and auto-segmented c-$Fos^+$ counts. **(N)** A high correlation (R = 0.89) was found between manual c-$Fos^+$ cell counts by Counter A and the number of auto-segmented objects. Each dot represents one ROI counted. **(O)** A high correlation (R = 0.93) was found between manual c-$Fos^+$ cell counts by Counter B and the number of auto-segmented objects. **(P)** There was high inter-rater correlation (R = 0.93) between manual c-$Fos^+$ cell counts. **(Q)** Averaged manual c-$Fos^+$ cell counts showed overall high correlation (R = 0.93) with auto-segmented counts. **(R)** Counts of all FP, FN, and TP cell counts when comparing auto-segmented c-$Fos^+$ counts and manual counts.

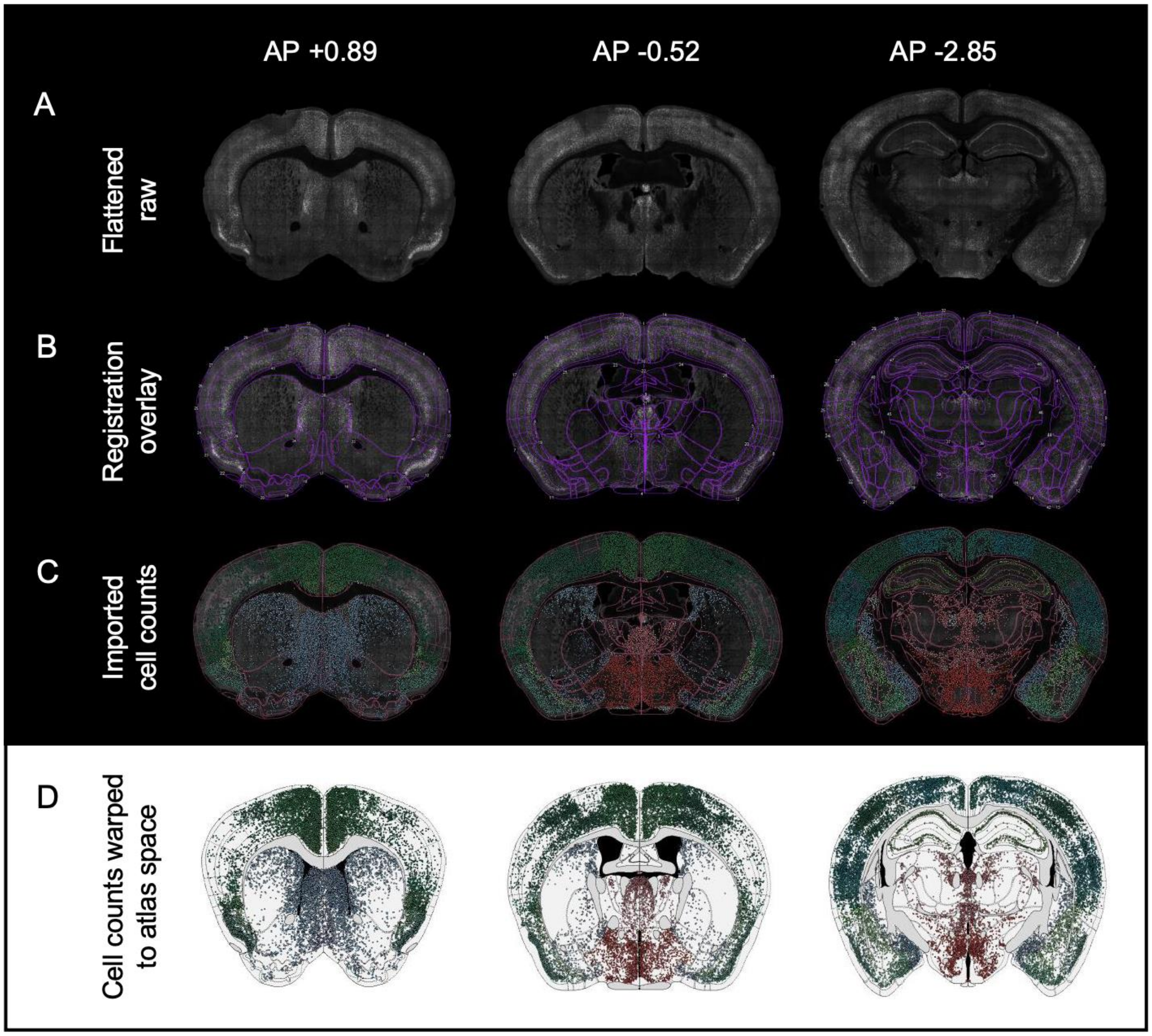


**Figure S2. Registration projecting segmented cell counts to a standard atlas space. (A)** Raw grayscale images of whole coronal sections with eYFP$^{+}$ and c-Fos$^{+}$ signals combined (Flattened raw). Corresponding coordinates of the best matching Allen Mouse Brain atlas plates (AP +0.89, left; AP -0.52, middle; AP -2.85, right) are displayed above. **(B)** The user-corrected registration overlay across the representative coronal sections. User-correction through an interactive console interface was performed by interfacing with the Wholebrain and SMART packages through SMARTTR. **(C)** Representative images of imported c-Fos automated cell counts onto the user-corrected registrations (image space). Cells are color coded based on region colors consistent with the Allen Mouse Brain Atlas. **(D)** Forward warp of the segmented c-Fos$^{+}$ counts from image space to a common shared stereotaxic coordinate space (atlas space) facilitates standardized integration of data across all mice.

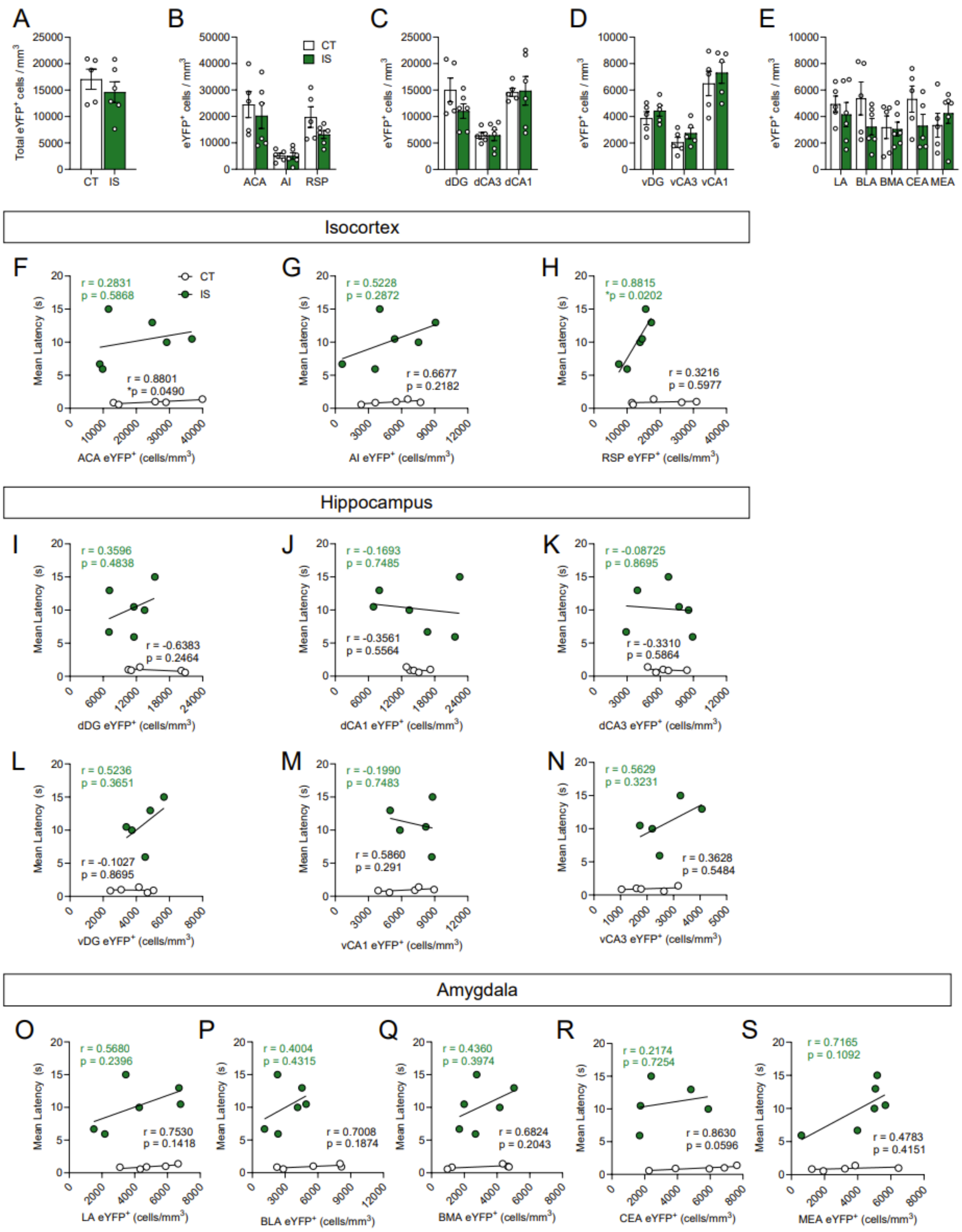


**Figure S3. Activity is not heightened globally or across targeted subregions following exposure to inescapable shock. (A)** No significant differences were found in total brain-wide volume normalized (cells/mm$^3$) counts of eYFP$^+$ cells between context trained (CT) or inescapable shock (IS) groups. Regional activation analysis of targeted **(B)** isocortical areas (ACA, AI, RSP), **(C)** dorsal hippocampal regions (dDG, dCA3, dCA1), **(D)** ventral hippocampal regions (vDG, vCA3, vCA1), and **(E)** amygdalar areas (LA, BLA, BMA, CEA) revealed no difference in neural activity levels between CT and IS groups. **(F-H)** Targeted isocortical analysis showed that eYFP$^+$ cell counts in the RSP and escape latency during later escapable shock testing were significantly correlated in the IS group. **(I-N)** No significant correlations were found between dorsal and ventral hippocampal region eYFP$^+$ cell counts and escape latency during later escapable shock testing. **(O-S)** No significant correlations were found between targeted amygdalar region eYFP cell counts and escape latency. Error bars represent ± SEM. Per brain region analyzed, n = 5 mice for the CT group and n = 5-6 for the IS group. ACA, anterior cingulate area; AI, agranular insula; RSP, retrosplenial area; dDG, dorsal dentate gyrus; dCA3, dorsal CA3; dCA1, dorsal CA1; vDG, ventral dentate gyrus; vCA3, ventral CA3; vCA1, ventral CA1; LA, lateral amygdalar nucleus; BLA, basolateral amygdalar nucleus; BMA, basomedial amygdalar nucleus; CEA, central amygdalar nucleus; MEA, medial amygdalar nucleus.

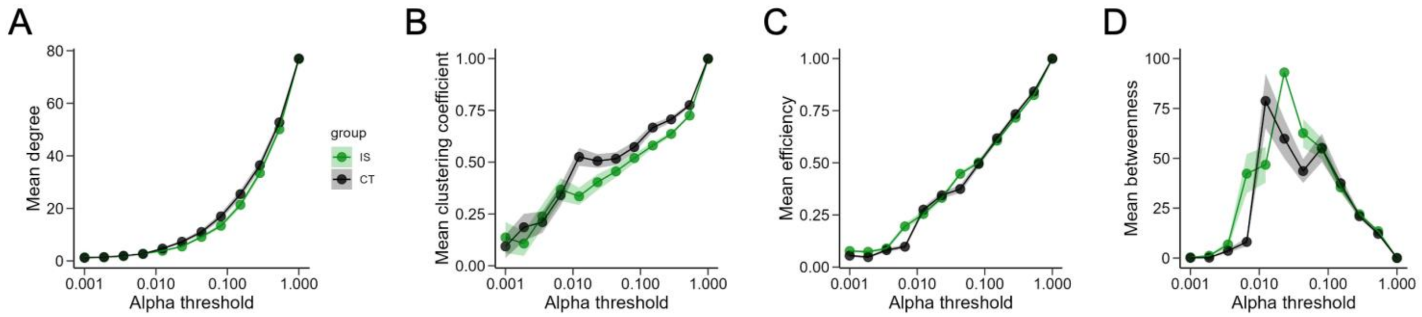


**Figure S4. Global network topology metrics during exposure to inescapable shock across a range of thresholds.** **(A)** Degree, **(B)** clustering coefficient, **(C)** global efficiency, and **(D)** mean betweenness centrality trajectories do not differ between CT and IS groups across a wide range of alpha thresholds.

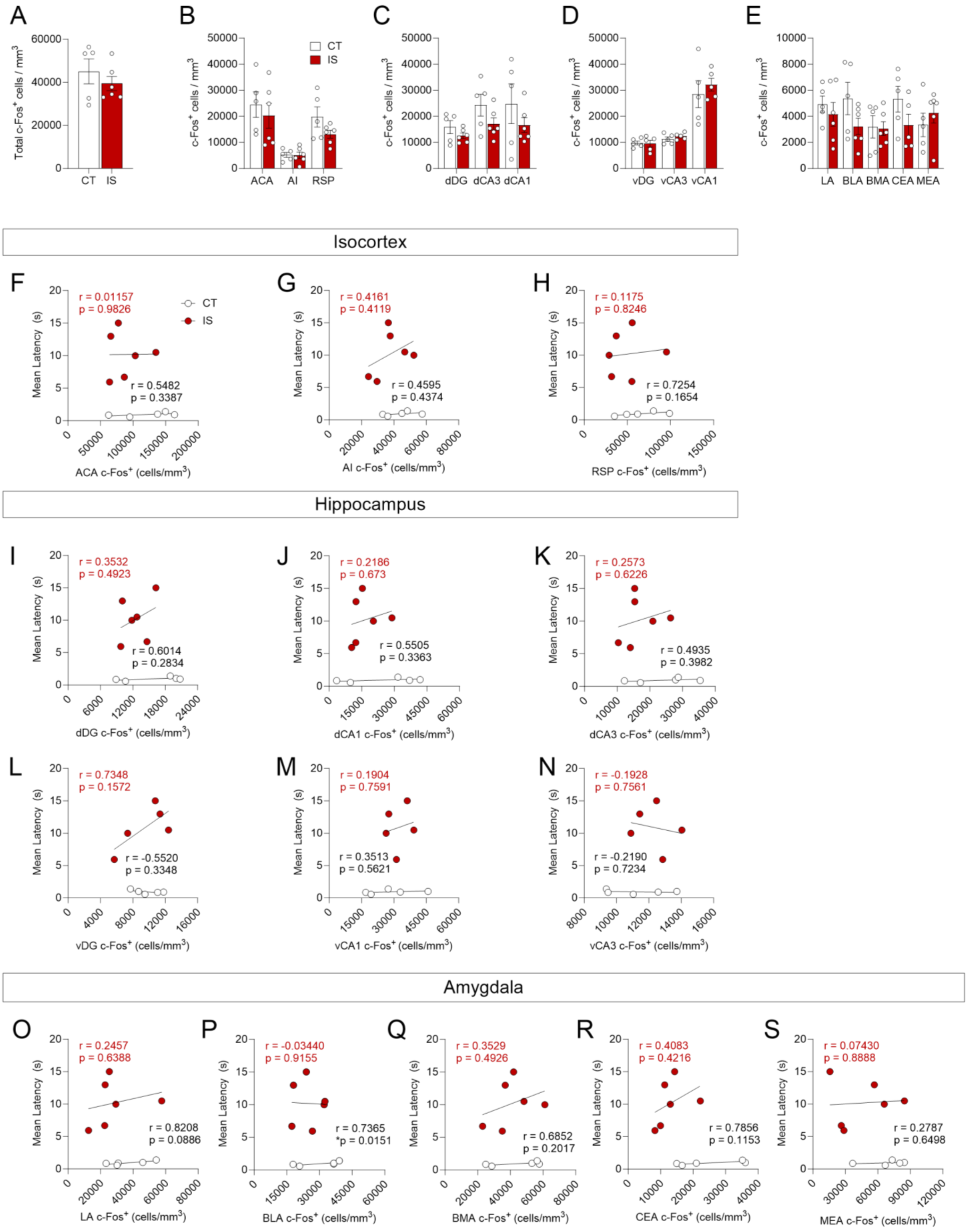


**Figure S5. Absolute activity of targeted isocortical, hippocampal, and amygdala regions are not**

**associated with learned helpless expression. (A)** No significant differences were found in total brain-wide volume normalized (cells/mm$^3$) counts of c-Fos$^+$ cells between context trained (CT) or inescapable shock (IS) groups. Regional activation analysis of targeted **(B)** isocortical areas (ACA, AI, RSP), **(C)** dorsal hippocampal regions (dDG, dCA3, dCA1), **(D)** ventral hippocampal regions (vDG, vCA3, vCA1), and **(E)** amygdalar areas (LA, BLA, BMA, CEA) revealed no difference in neural activity levels between CT and IS groups. **(F-H)** c-Fos$^+$ cell counts in targeted isocortical regions do not correlate with escape latency during later escapable shock. **(I-N)** No significant correlations were found between dorsal and ventral hippocampal region c-Fos$^+$ cell counts and escape latency during later escapable shock testing. **(O-S)** No significant correlations were found between targeted amygdalar region c-Fos$^+$ cell counts and escape latency. Error bars represent ± SEM. Per brain region analyzed, n = 5 mice for the CT group and n = 5-6 for the IS group. ACA, anterior cingulate area; AI, agranular insula; RSP, retrosplenial area; dDG, dorsal dentate gyrus; dCA3, dorsal CA3; dCA1, dorsal CA1; vDG, ventral dentate gyrus; vCA3, ventral CA3; vCA1, ventral CA1; LA, lateral amygdalar nucleus; BLA, basolateral amygdalar nucleus; BMA, basomedial amygdalar nucleus; CEA, central amygdalar nucleus; MEA, medial amygdalar nucleus.

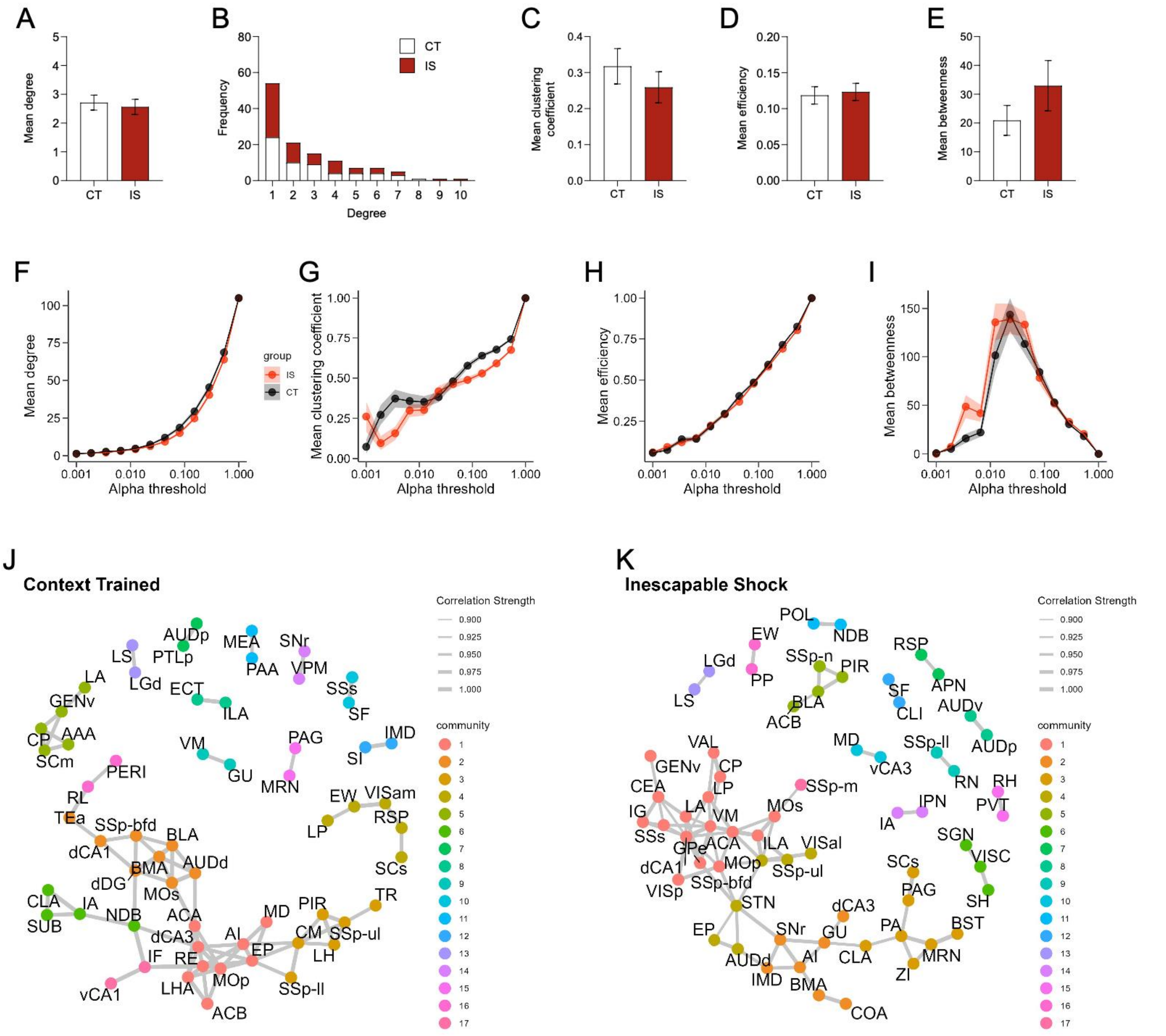


**Figure S6. Properties of networks constructed from brain-wide c-Fos+ regional co-expression.**

**(A)** Average degree does not differ between CT and IS groups. **(B)** Degree frequency distribution shows a right-tailed distribution for both groups. **(C-E)** Mean clustering coefficient, global efficiency, and mean betweenness centrality does not differ between groups. **(F-I)** Trajectories of global c-Fos$^{+}$ network topology metrics of mean degree, mean clustering coefficient, mean efficiency, and mean betweenness centrality are calculated based off a wide range of possible significance thresholds. Exploratory community detection analysis applied to **(J)** Context trained (CT) and **(K)** inescapable shock (IS) c-Fos$^{+}$ networks based on calculation of the leading non-negative eigenvector of the modularity matrix. Gray lines indicate retained network connections between nodes. Nodes belonging to the same detected communities are colored identically. Error shading represents ± SEM; n = 5 mice for the CT group and n = 6 for the IS group.

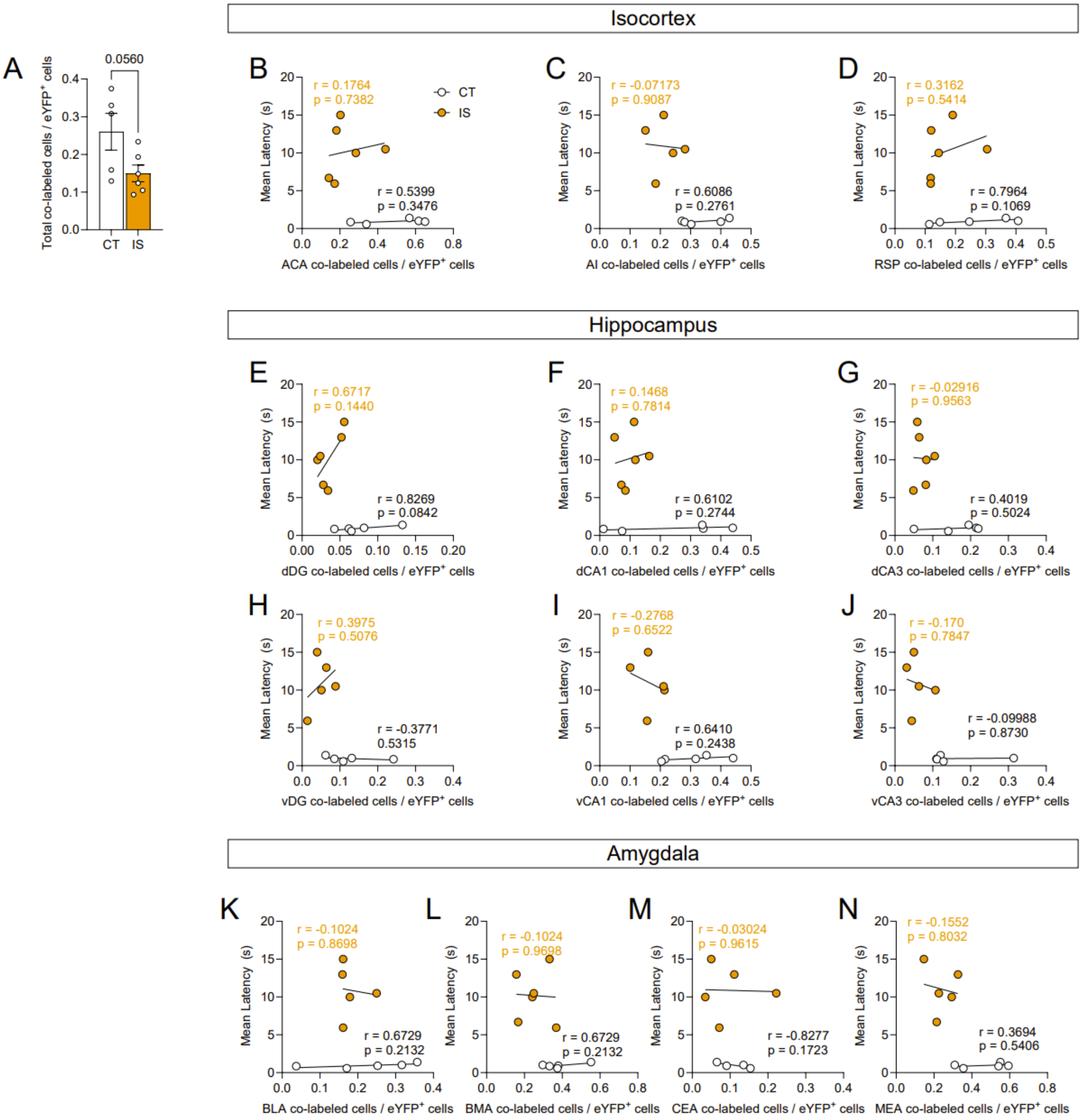


**Figure S7. Global reactivation activity during escapable shock is dampened in mice previous exposed to inescapable shock. (A)** There is a trending global decrease of proportion of reactivated cells (co-labeled cells / eYFP$^{+}$ cells) in inescapable shock (IS) mice compared to context trained (CT) mice. Reactivation activity of targeted isocortical areas, such as the **(B)** ACA, **(C)** AI, and **(D)** RSP, is not correlated to escape latency in IS or CT mice. Reactivation activity of targeted dorsal hippocampal regions, such as the **(E)** dDG, **(F)** dCA1, **(G)** and dCA3, and of targeted ventral hippocampal regions, such as the **(H)** vDG, **(I)** vCA1, **(J)** and vCA3, is not correlated to escape latency in IS or CT mice. Reactivation activity of targeted amygdalar areas, such as the **(K)** BLA, **(L)** BMA, **(M)** CEA, **(N)** and MEA, is not correlated to escape latency in IS or CT mice. Error bars represent ± SEM. Per brain region analyzed, n = 4-5 mice for the CT group and n = 5-6 for the IS group. ACA, anterior cingulate area; AI, agranular insula; RSP, retrosplenial area; dDG, dorsal dentate gyrus; dCA3, dorsal CA3; dCA1, dorsal CA1; vDG , ventral dentate gyrus; vCA3, ventral CA3; vCA1, ventral CA1; LA, lateral amygdalar nucleus; BLA, basolateral amygdalar nucleus; BMA, basomedial amygdalar nucleus; CEA, central amygdalar nucleus; MEA, medial amygdalar nucleus.

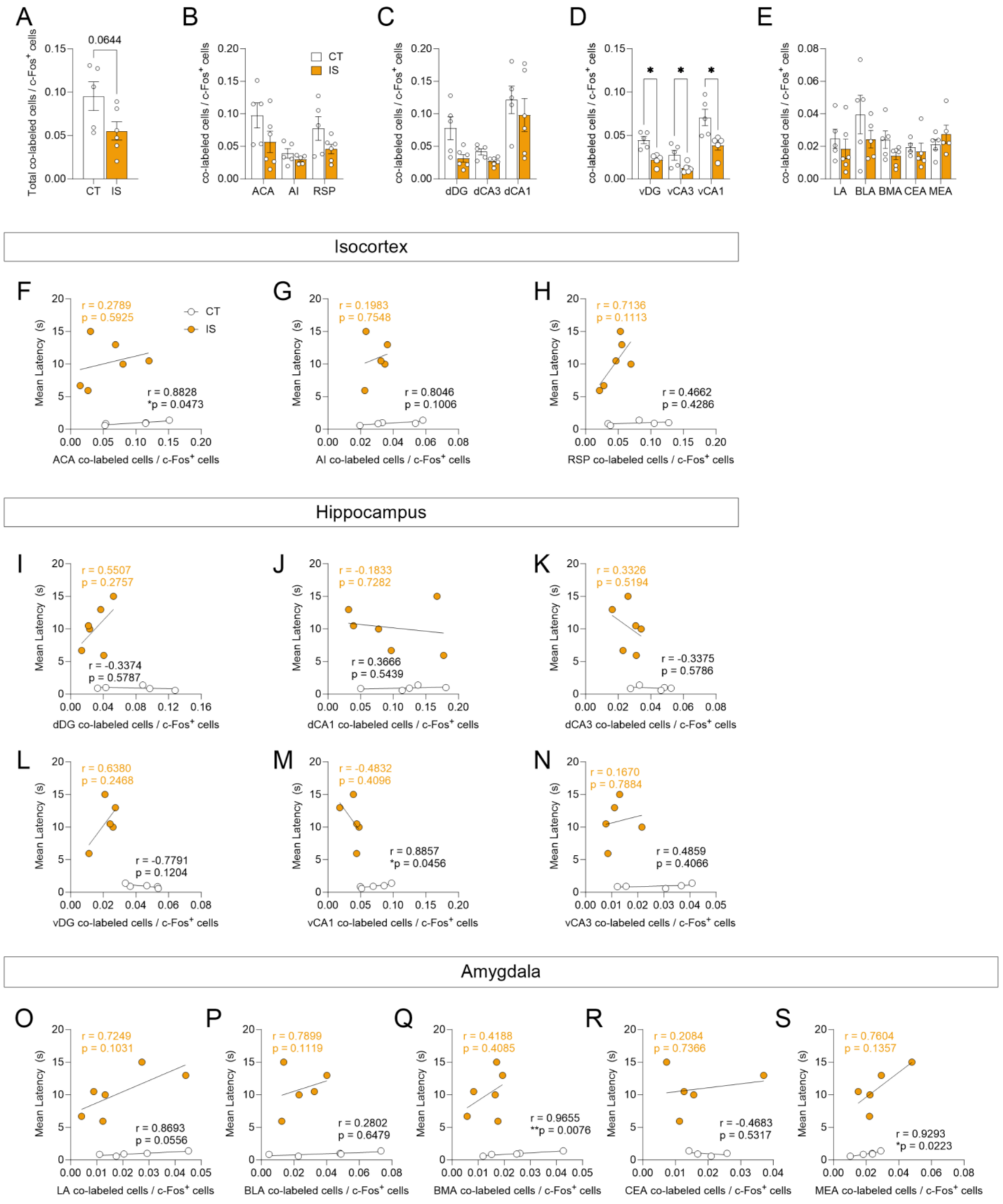

**Figure S8. Reactivated activity proportions are decreased in targeted ventral hippocampal regions following inescapable shock. (A)** There is a trending decrease in global co-labeled / c-Fos$^+$ cell proportions between context trained (CT) or inescapable shock (IS) groups. Regional activation analysis of targeted **(B)**

isocortical areas (ACA, AI, RSP), **(C)** dorsal hippocampal regions (dDG, dCA3, dCA1), **(D)** ventral hippocampal regions (vDG, vCA3, vCA1), and **(E)** amygdalar areas (LA, BLA, BMA, CEA) reveal that vDG, vCA3, and vCA1 activity are significantly decreased in IS mice compared to CT mice. Correlation analysis of co-labeled / c-Fos+ activity of targeted isocortical areas, such as the **(F)** ACA, **(G)** AI, and **(H)** RSP, reveal that ACA activity is significantly correlated with crossing latency in the CT group. Analysis of targeted dorsal and ventral hippocampal regions, such as the **(I)** dDG, **(J)** dCA1, **(K)** dCA3, **(L)** vDG, **(M)** vCA1, **(N)** and vCA3, reveal that reveal that vCA1 co-labeled / c-Fos+ is significantly correlated with crossing latency in the CT group. Analysis of targeted amygdalar regions, such as **(O)** LA, **(P)** BLA, **(Q)** BMA, **(R)** CEA, **(S)** MEA reveal that reveal that co-labeled / c-Fos+ activity in the BMA and MEA is significantly correlated with crossing latency in the CT group. Error bars represent ± SEM. Per brain region, n = 4-5 mice for the CT group and n = 5-6 for the IS group. ACA, anterior cingulate area; AI, agranular insula; RSP, retrosplenial area; dDG, dorsal dentate gyrus; dCA3, dorsal CA3; dCA1, dorsal CA1; vDG, ventral dentate gyrus; vCA3, ventral CA3; vCA1, ventral CA1; LA, lateral amygdalar nucleus; BLA, basolateral amygdalar nucleus; BMA, basomedial amygdalar nucleus; CEA, central amygdalar nucleus; MEA, medial amygdalar nucleus.

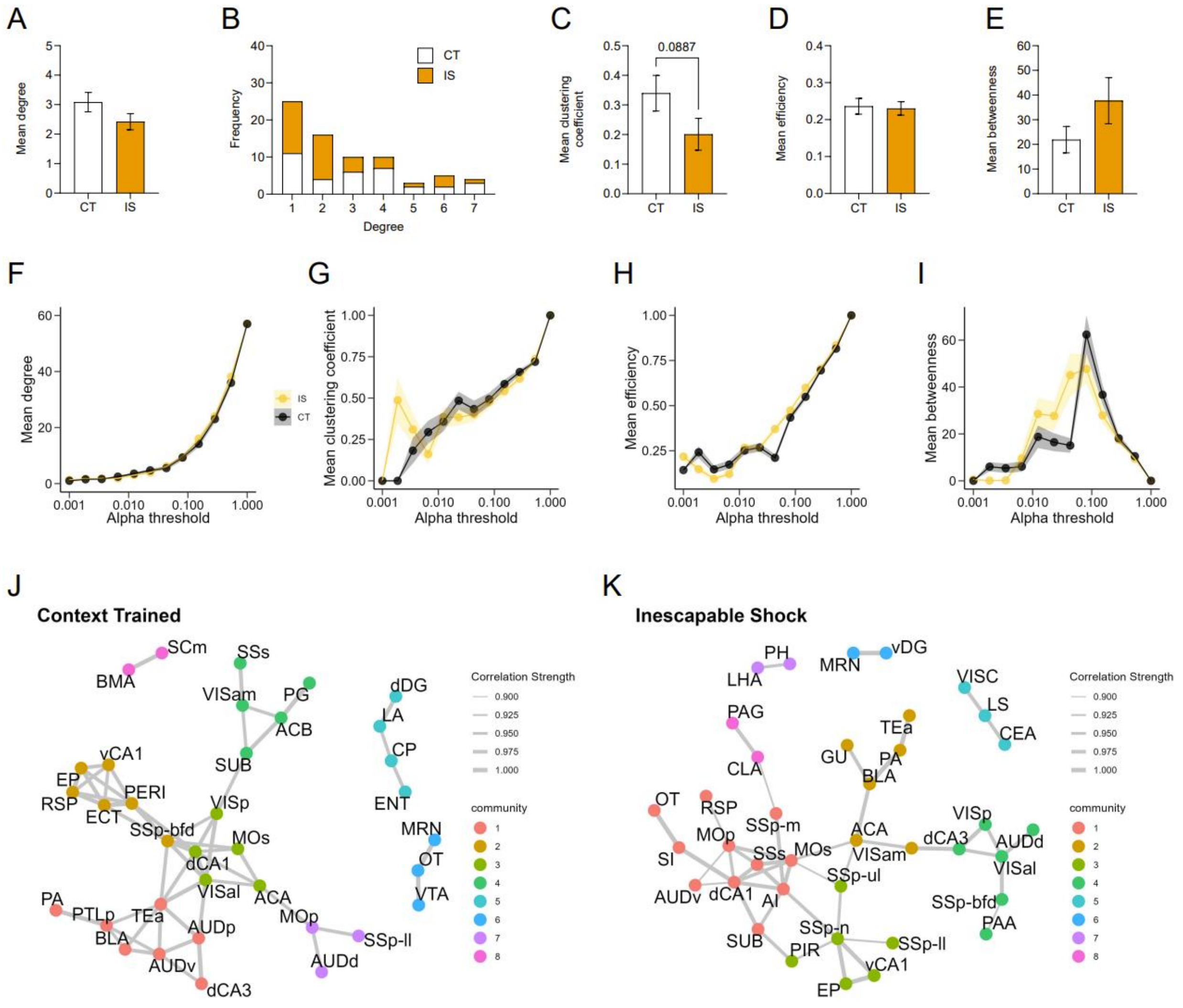


**Figure S9. Properties of networks constructed from functional connections between reactivated ensembles (co-labeled / eYFP⁺ cells).** Global topology characteristics of CT and IS reactivation networks were calculated after thresholding for important regional correlations using an $r > 0.9$ and $p < 0.01$. **(A)** Average degree does not differ between groups. **(B)** Degree frequency distribution shows a right-tailed distribution for both groups. **(C-E)** Mean clustering coefficient, global efficiency, and mean betweenness centrality does not differ between groups. Trajectories of global network topology metrics, including **(F)** mean degree, **(G)** mean clustering coefficient, **(H)** mean efficiency, and **(I)** mean betweenness, were calculated based off a wide range of possible significance thresholds. Exploratory community detection analysis applied to **(J)** CT and **(K)** IS reactivation networks based on calculation of the leading non-negative eigenvector of the modularity matrix. Gray lines indicate retained network connections between nodes. Nodes belonging to the same detected communities are identically colored. Error bars represent ± SEM; n = 5 mice for the CT group and n = 6 for the IS group.

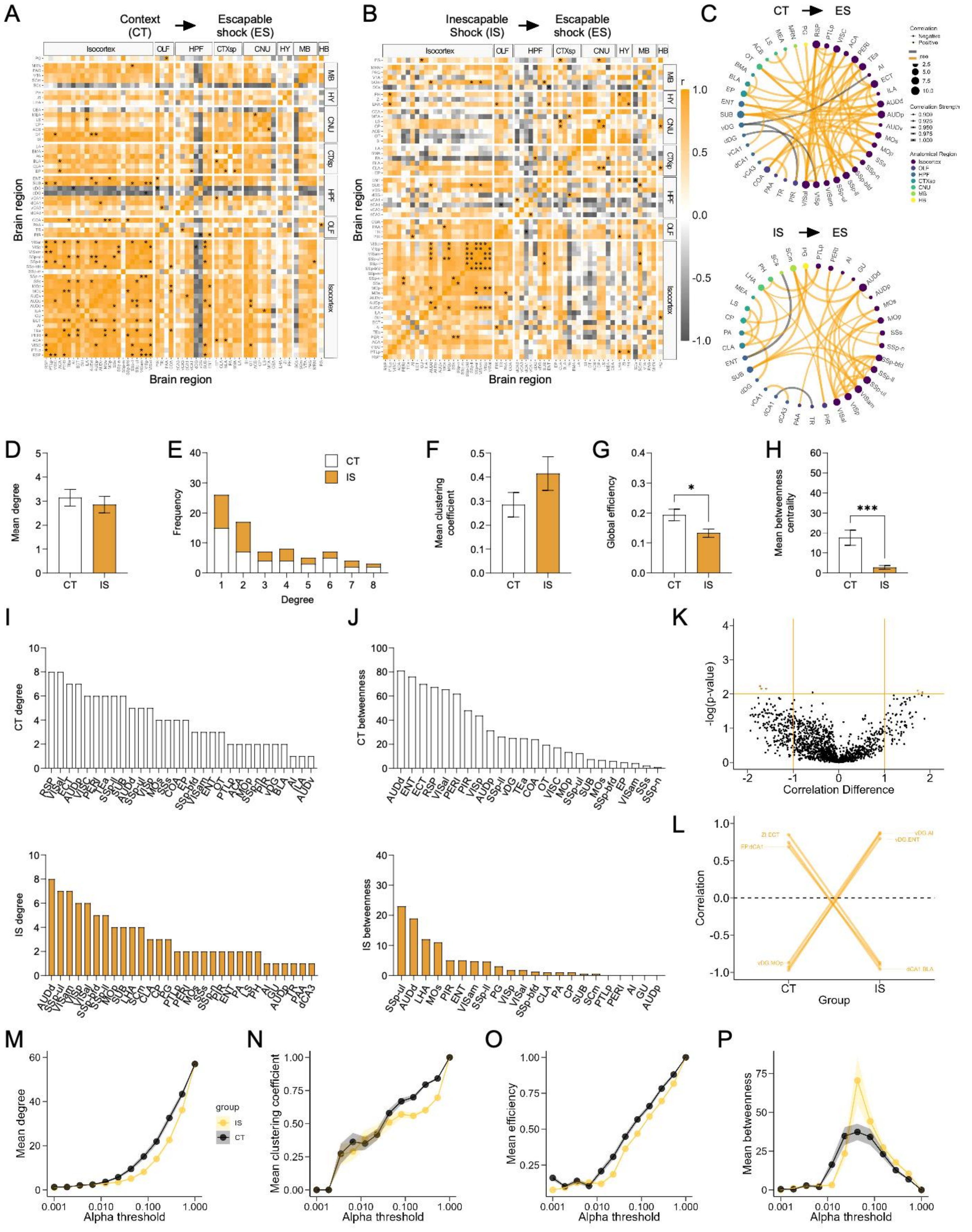

**Figure S10. Supplementary network level analysis of co-labeled / c-Fos$^+$ activity reveals high sensory functional connectivity. (A-B)** Regional cross correlation heatmaps of colabel / c-Fos$^+$ proportions in context trained (CT) and learned helpless (IS) mice. Significant values are $p < 0.01$. In the IS heatmaps, an especially high co-activation amongst somatosensory regions, visual, and auditory regions is observed. **(C)** Functional networks constructed after thresholding for the strongest and most significantly correlated or anti-correlated connections ($r > 0.9$, $p < 0.01$). **(D)** Average degree centrality does not differ between IS and CT groups. **(E)** Degree frequency distributions are right-tailed. **(F)** Mean clustering coefficient does not differ between the CT and IS networks. **(G-H)** Mean global efficiency and betweenness centrality are significantly lower in IS networks, likely due to overall lower interconnectivity amongst regions. (**I**) The top node degree values in descending order for the CT (white, top) and IS (yellow, bottom) networks indicate which regions are most highly connected. **(J)** The top node betweenness values in descending order for the CT (white, top) and IS (yellow, bottom) networks. **(K)** Volcano plot of the Pearson correlation differences ($r_{IS} – r_{CT}$) for all individual regional connections against their p-values calculated from a permutation analysis. Points intersecting or within the upper left or right quadrant represent the regional relationships with the greatest change (|correlation difference| > 1), that were most significant ($p < 0.01$). **(L)** A parallel coordinate plot highlighting individual significantly changed regional correlations between groups, as well as the direction of their change. **(M-P)** Trajectories of global c-Fos$^+$ network topology metrics of mean degree, mean efficiency, mean betweenness centrality, and mean clustering coefficients are calculated based off a wide range of possible significance thresholds. ***$p < 0.001$. Error bars represent ± SEM; n = 5 mice for the CT group and n = 6 for the IS group.

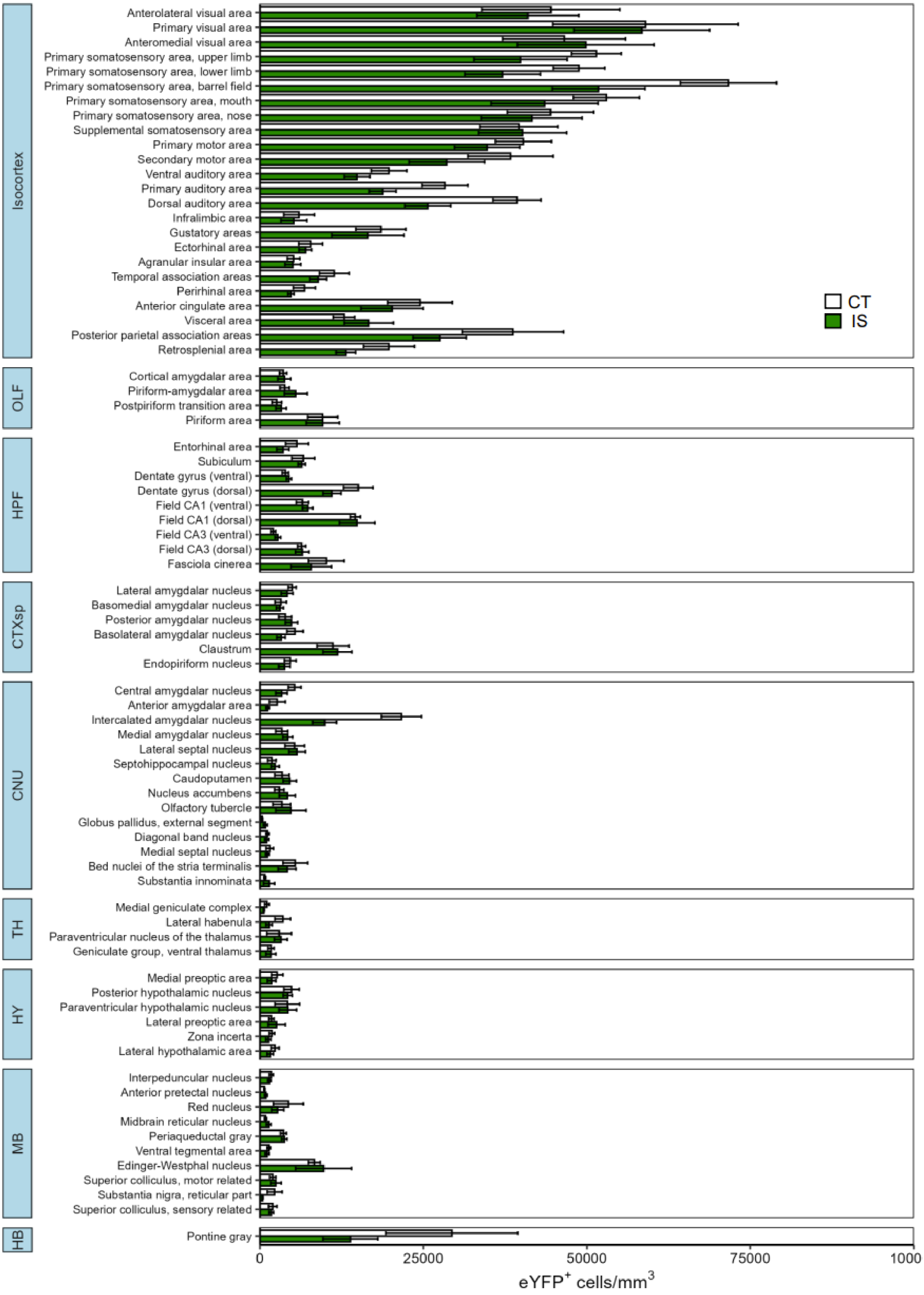
Isocortex
Anterolateral visual area
Primary visual area
Anteromedial visual area
Primary somatosensory area, upper limb
Primary somatosensory area, lower limb
Primary somatosensory area, barrel field
Primary somatosensory area, mouth
Primary somatosensory area, nose
Supplemental somatosensory area
Primary motor area
Secondary motor area
Ventral auditory area
Primary auditory area
Dorsal auditory area
Infralimbic area
Gustatory areas
Ectorhinal area
Agranular insular area
Temporal association areas
Perirhinal area
Anterior cingulate area
Visceral area
Posterior parietal association areas
Retrosplenial area
CT
IS
OLF
Cortical amygdalar area
Piriform-amygdalar area
Postpiriform transition area
Piriform area
HPF
Entorhinal area
Subiculum
Dentate gyrus (ventral)
Dentate gyrus (dorsal)
Field CA1 (ventral)
Field CA1 (dorsal)
Field CA3 (ventral)
Field CA3 (dorsal)
Fasciola cinerea
CTXsp
Lateral amygdalar nucleus
Basomedial amygdalar nucleus
Posterior amygdalar nucleus
Basolateral amygdalar nucleus
Claustrum
Endopiriform nucleus
CNU
Central amygdalar nucleus
Anterior amygdalar area
Intercalated amygdalar nucleus
Medial amygdalar nucleus
Lateral septal nucleus
Septohippocampal nucleus
Caudoputamen
Nucleus accumbens
Olfactory tubercle
Globus pallidus, external segment
Diagonal band nucleus
Medial septal nucleus
Bed nuclei of the stria terminalis
Substantia innominata
TH
Medial geniculate complex
Lateral habenula
Paraventricular nucleus of the thalamus
Geniculate group, ventral thalamus
HY
Medial preoptic area
Posterior hypothalamic nucleus
Paraventricular hypothalamic nucleus
Lateral preoptic area
Zona incerta
Lateral hypothalamic area
MB
Interpeduncular nucleus
Anterior pretectal nucleus
Red nucleus
Midbrain reticular nucleus
Periaqueductal gray
Ventral tegmental area
Edinger-Westphal nucleus
Superior colliculus, motor related
Substantia nigra, reticular part
Superior colliculus, sensory related
HB
Pontine gray
0
25000
50000
75000
1000
eYFP+ cells/mm3

l., 20

**Figure S11. Differential expression of eYFP activity across all mapped regions.** Volume normalized (cells/mm$^3$) counts of eYFP$^+$ cells between context trained (CT, white bars) or inescapable shock (IS, green bars) mice across all regions mapped. Subregion activity expression patterns are organized by parent anatomical divisions. Error bars represent ± SEM. OLF, olfactory areas; HPF, hippocampal formation; CTXsp, cortical subplate; CNU, cerebral nuclei; TH, thalamus; HY, hypothalamus; MB, midbrain; HB, hindbrain. Error bars represent ± SEM; n = 5 mice for the CT group and n = 6 for the IS group.

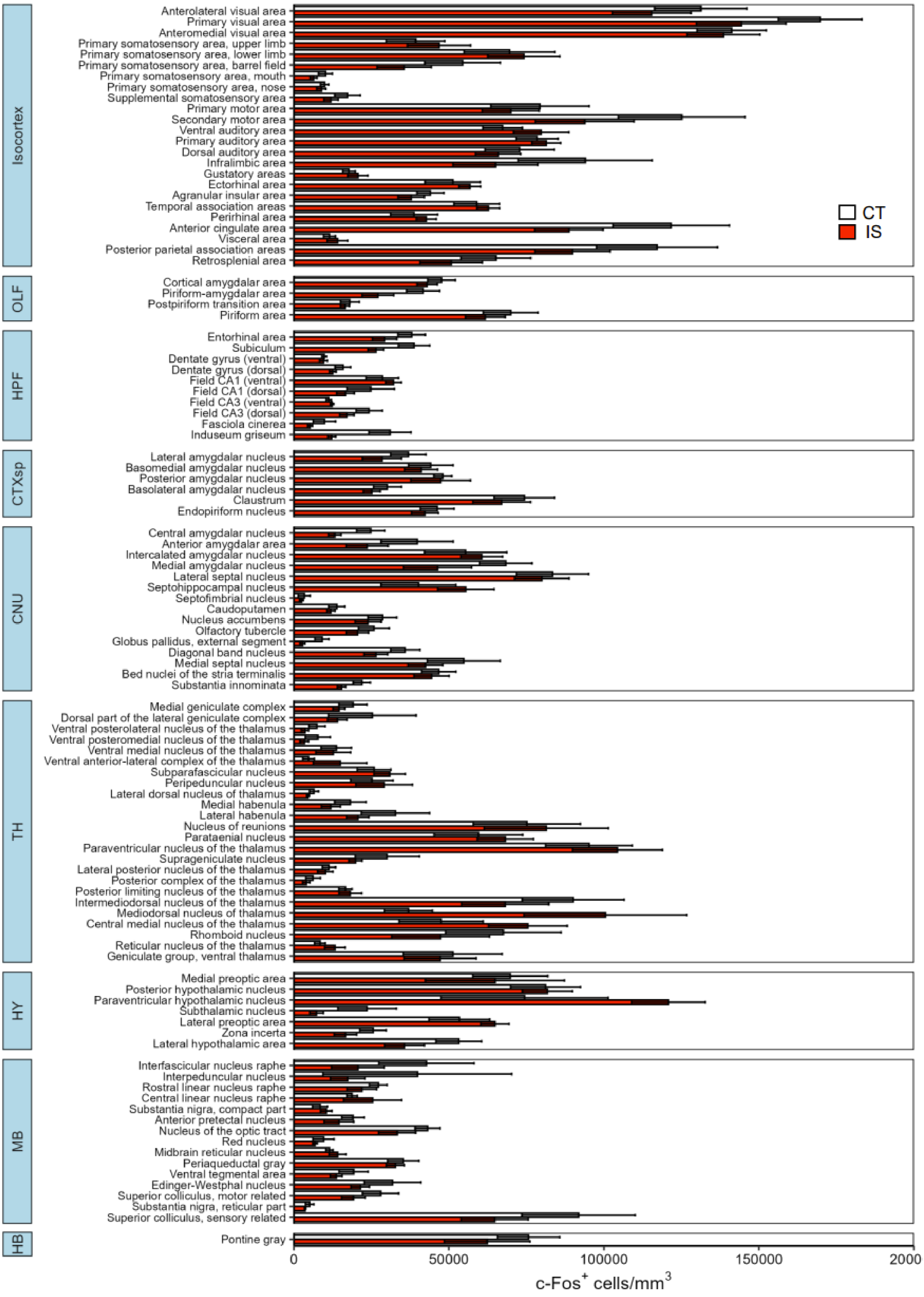
Isocortex
Anterolateral visual area
Primary visual area
Anteromedial visual area
Primary somatosensory area, upper limb
Primary somatosensory area, lower limb
Primary somatosensory area, barrel field
Primary somatosensory area, mouth
Primary somatosensory area, nose
Supplemental somatosensory area
Primary motor area
Secondary motor area
Ventral auditory area
Primary auditory area
Dorsal auditory area
Infralimbic area
Gustatory areas
Ectorhinal area
Agranular insular area
Temporal association areas
Perirhinal area
Anterior cingulate area
Visceral area
Posterior parietal association areas
Retrosplenial area
OLF
Cortical amygdalar area
Piriform-amygdalar area
Postpiriform transition area
Piriform area
HPF
Entorhinal area
Subiculum
Dentate gyrus (ventral)
Dentate gyrus (dorsal)
Field CA1 (ventral)
Field CA1 (dorsal)
Field CA3 (ventral)
Field CA3 (dorsal)
Fasciola cinerea
Induseum griseum
CTXsp
Lateral amygdalar nucleus
Basomedial amygdalar nucleus
Posterior amygdalar nucleus
Basolateral amygdalar nucleus
Claustrum
Endopiriform nucleus
CNU
Central amygdalar nucleus
Anterior amygdalar area
Intercalated amygdalar nucleus
Medial amygdalar nucleus
Lateral septal nucleus
Septohippocampal nucleus
Septofimbrial nucleus
Caudoputamen
Nucleus accumbens
Olfactory tubercle
Globus pallidus, external segment
Diagonal band nucleus
Medial septal nucleus
Bed nuclei of the stria terminalis
Substantia innominata
TH
Medial geniculate complex
Dorsal part of the lateral geniculate complex
Ventral posterolateral nucleus of the thalamus
Ventral posteromedial nucleus of the thalamus
Ventral medial nucleus of the thalamus
Ventral anterior-lateral complex of the thalamus
Subparafascicular nucleus
Peripeduncular nucleus
Lateral dorsal nucleus of thalamus
Medial habenula
Lateral habenula
Nucleus of reunions
Parataenial nucleus
Paraventricular nucleus of the thalamus
Suprageniculate nucleus
Lateral posterior nucleus of the thalamus
Posterior complex of the thalamus
Posterior limiting nucleus of the thalamus
Intermediodorsal nucleus of the thalamus
Mediodorsal nucleus of thalamus
Central medial nucleus of the thalamus
Rhomboid nucleus
Reticular nucleus of the thalamus
Geniculate group, ventral thalamus
HY
Medial preoptic area
Posterior hypothalamic nucleus
Paraventricular hypothalamic nucleus
Subthalamic nucleus
Lateral preoptic area
Zona incerta
Lateral hypothalamic area
MB
Interfascicular nucleus raphe
Interpeduncular nucleus
Rostral linear nucleus raphe
Central linear nucleus raphe
Substantia nigra, compact part
Anterior pretectal nucleus
Nucleus of the optic tract
Red nucleus
Midbrain reticular nucleus
Periaqueductal gray
Ventral tegmental area
Edinger-Westphal nucleus
Superior colliculus, motor related
Substantia nigra, reticular part
Superior colliculus, sensory related
HB
Pontine gray
CT
IS
0
50000
100000
150000
2000
c-Fos+ cells/mm3

**Figure S12. Differential expression of c-Fos activity across all mapped regions.** Volume normalized (cells/$mm^3$) counts of c-$Fos^+$ cells between context trained (CT, white bars) or inescapable shock (IS, red bars) mice across all regions mapped. Subregion activity expression patterns are organized by parent anatomical divisions. Error bars represent ± SEM. OLF, olfactory areas; HPF, hippocampal formation; CTXsp, cortical subplate; CNU, cerebral nuclei; TH, thalamus; HY, hypothalamus; MB, midbrain; HB, hindbrain. Error bars represent ± SEM; n = 5 mice for the CT group and n = 6 for the IS group.

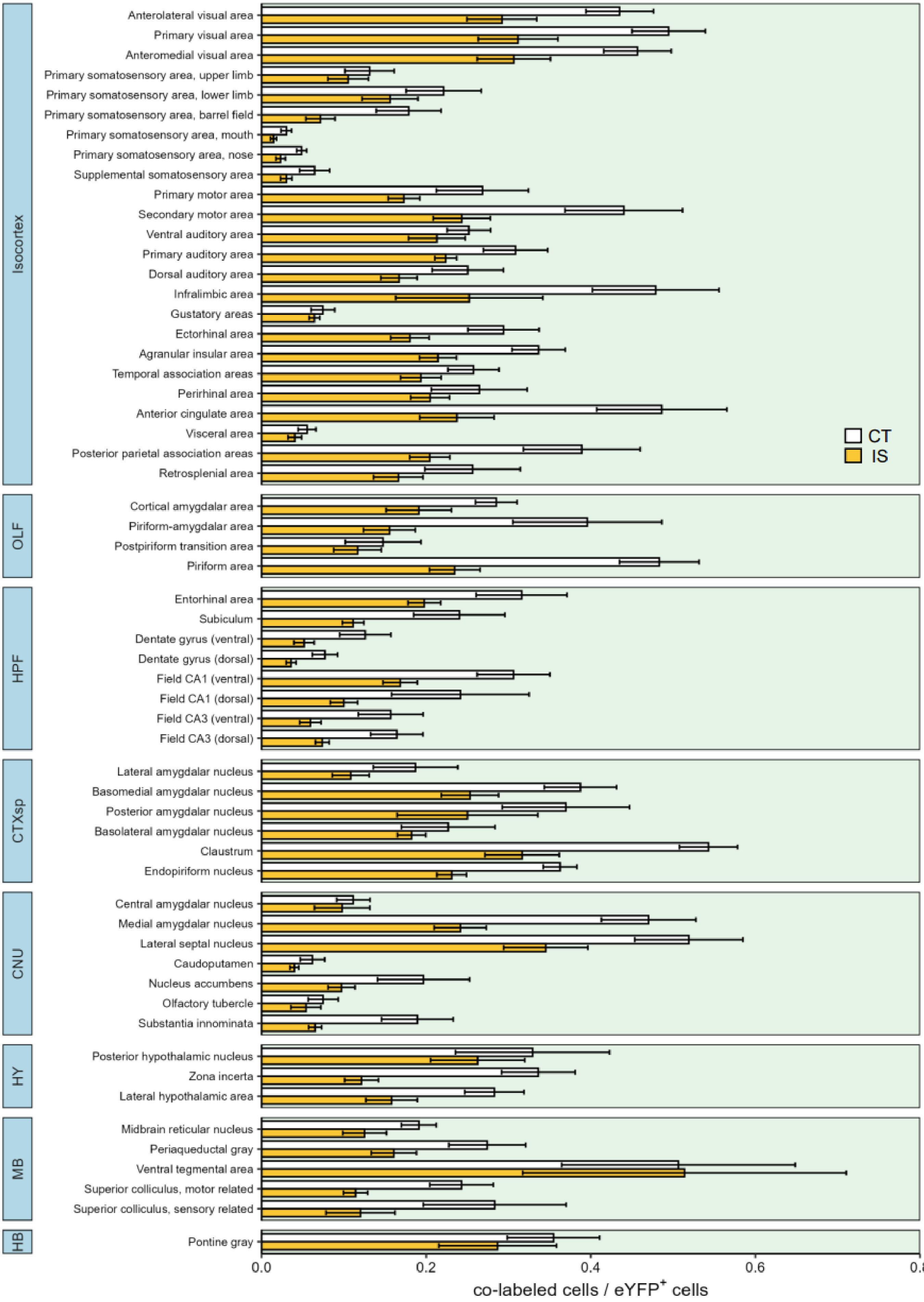
Isocortex
Anterolateral visual area
Primary visual area
Anteromedial visual area
Primary somatosensory area, upper limb
Primary somatosensory area, lower limb
Primary somatosensory area, barrel field
Primary somatosensory area, mouth
Primary somatosensory area, nose
Supplemental somatosensory area
Primary motor area
Secondary motor area
Ventral auditory area
Primary auditory area
Dorsal auditory area
Infralimbic area
Gustatory areas
Ectorhinal area
Agranular insular area
Temporal association areas
Perirhinal area
Anterior cingulate area
Visceral area
Posterior parietal association areas
Retrosplenial area
OLF
Cortical amygdalar area
Piriform-amygdalar area
Postpiriform transition area
Piriform area
HPF
Entorhinal area
Subiculum
Dentate gyrus (ventral)
Dentate gyrus (dorsal)
Field CA1 (ventral)
Field CA1 (dorsal)
Field CA3 (ventral)
Field CA3 (dorsal)
CTXsp
Lateral amygdalar nucleus
Basomedial amygdalar nucleus
Posterior amygdalar nucleus
Basolateral amygdalar nucleus
Claustrum
Endopiriform nucleus
CNU
Central amygdalar nucleus
Medial amygdalar nucleus
Lateral septal nucleus
Caudoputamen
Nucleus accumbens
Olfactory tubercle
Substantia innominata
HY
Posterior hypothalamic nucleus
Zona incerta
Lateral hypothalamic area
MB
Midbrain reticular nucleus
Periaqueductal gray
Ventral tegmental area
Superior colliculus, motor related
Superior colliculus, sensory related
HB
Pontine gray
CT
IS
0.0
0.2
0.4
0.6
0.8
co-labeled cells / eYFP$^+$ cells

**Figure S13. Differential reactivation proportions across all mapped regions.** Reactivation activity (colabelled cells / eYFP$^+$ cells) between context trained (CT, white bars) or inescapable shock (IS, yellow bars) mice across all regions mapped. Subregion activity expression patterns are organized by parent anatomical divisions. Error bars represent ± SEM. OLF, olfactory areas; HPF, hippocampal formation; CTXsp, cortical subplate; CNU, cerebral nuclei; TH, thalamus; HY, hypothalamus; MB, midbrain; HB, hindbrain. Error bars represent ± SEM; n = 5 mice for the CT group and n = 6 for the IS group.

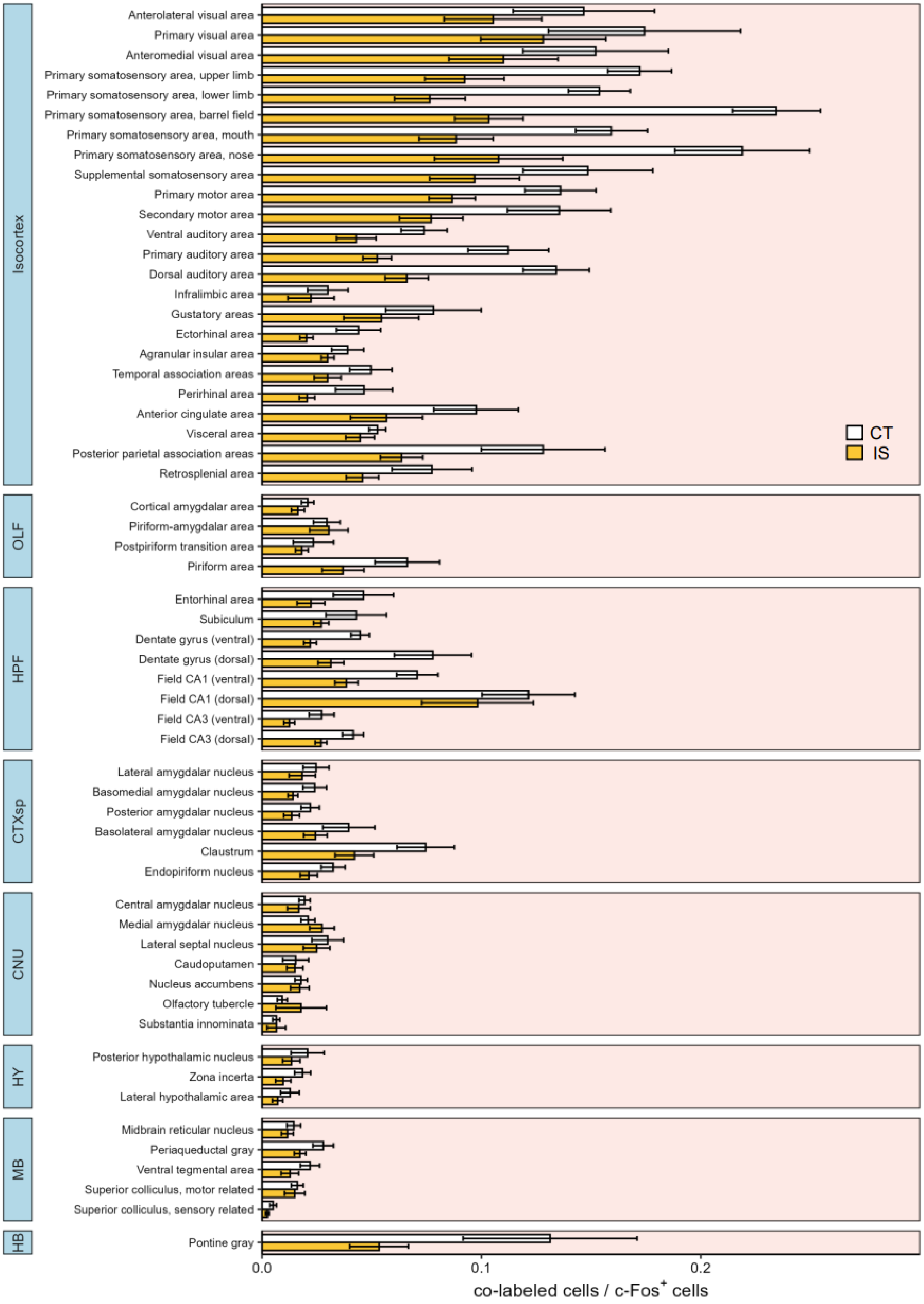
Isocortex
Anterolateral visual area
Primary visual area
Anteromedial visual area
Primary somatosensory area, upper limb
Primary somatosensory area, lower limb
Primary somatosensory area, barrel field
Primary somatosensory area, mouth
Primary somatosensory area, nose
Supplemental somatosensory area
Primary motor area
Secondary motor area
Ventral auditory area
Primary auditory area
Dorsal auditory area
Infralimbic area
Gustatory areas
Ectorhinal area
Agranular insular area
Temporal association areas
Perirhinal area
Anterior cingulate area
Visceral area
Posterior parietal association areas
Retrosplenial area
CT
IS
OLF
Cortical amygdalar area
Piriform-amygdalar area
Postpiriform transition area
Piriform area
HPF
Entorhinal area
Subiculum
Dentate gyrus (ventral)
Dentate gyrus (dorsal)
Field CA1 (ventral)
Field CA1 (dorsal)
Field CA3 (ventral)
Field CA3 (dorsal)
CTXsp
Lateral amygdalar nucleus
Basomedial amygdalar nucleus
Posterior amygdalar nucleus
Basolateral amygdalar nucleus
Claustrum
Endopiriform nucleus
CNU
Central amygdalar nucleus
Medial amygdalar nucleus
Lateral septal nucleus
Caudoputamen
Nucleus accumbens
Olfactory tubercle
Substantia innominata
HY
Posterior hypothalamic nucleus
Zona incerta
Lateral hypothalamic area
MB
Midbrain reticular nucleus
Periaqueductal gray
Ventral tegmental area
Superior colliculus, motor related
Superior colliculus, sensory related
HB
Pontine gray
0.0
0.1
0.2
co-labeled cells / c-Fos+ cells

**Figure S14. Differential co-labeled / c-Fos$^+$ proportions across all mapped regions.** Co-labeled/c-Fos$^+$ proportions between context trained (CT, white bars) or inescapable shock (IS, yellow bars) mice across all regions mapped. Subregion activity expression patterns are organized by parent anatomical divisions. Error bars represent ± SEM. OLF, olfactory areas; HPF, hippocampal formation; CTXsp, cortical subplate; CNU, cerebral nuclei; TH, thalamus; HY, hypothalamus; MB, midbrain; HB, hindbrain. Error bars represent ± SEM; n = 5 mice for the CT group and n = 6 for the IS group.

### SUPPLEMENTAL TABLE LEGENDS

**Table S1. Key resources summary.**

**Table S2. Statistical analysis summary for all behavioral tests.**

**Table S3. Comprehensive list of acronyms used.**

**Table S4. Statistical analysis summary for targeted brain region analyses.**

**Table S5. Statistical analysis summary for eYFP activity across all mapped regions.** False discovery rate correction of p-values can be found under the p.adj column.

**Table S6. Statistical analysis summary for c-Fos activity across all mapped regions.** False discovery rate correction of p-values can be found under the p.adj column.

**Table S7. Statistical analysis summary for reactivation proportion (co-labeled cells / eYFP$^+$ cells) across all mapped regions.** False discovery rate correction of p-values can be found under the p.adj column.

**Table S8. Statistical analysis summary for proportion of reactivation activity (co-labeled cells / c-Fos$^+$ cells) across all mapped regions.** False discovery rate correction of p-values can be found under the p.adj column.
